## Supplementary Material for "Molecular basis of neurodegeneration in a mouse model of *Polr3*-related disease"

### Contents

**Supplementary Fig. S1.** Analysis of recombination frequency in Polr3a-tamKI mice.

**Supplementary Fig. S2.** Cerebellar histology and glucose homeostasis of Polr3a-tamKI mice.

**Supplementary Fig. S3.** Behavioral spectrometer analysis of WT and Polr3a-tamKI mice.

**Supplementary Fig. S4.** Gene expression in WT and Polr3a-tamKI (KI) cerebra at P75.

**Supplementary Fig. S5.** Pol III and Pol II transcript levels in various tissues from WT and Polr3a-tamKI (KI) mice at P75.

**Supplementary Fig. S6.** Gene expression and Iba1 staining in adolescent WT and Polr3a-tamKI (KI) cerebra.

**Supplementary Fig. S7.** Gene expression in WT and Polr3a-tamKI (KI) cerebella at P42.

**Supplementary references**

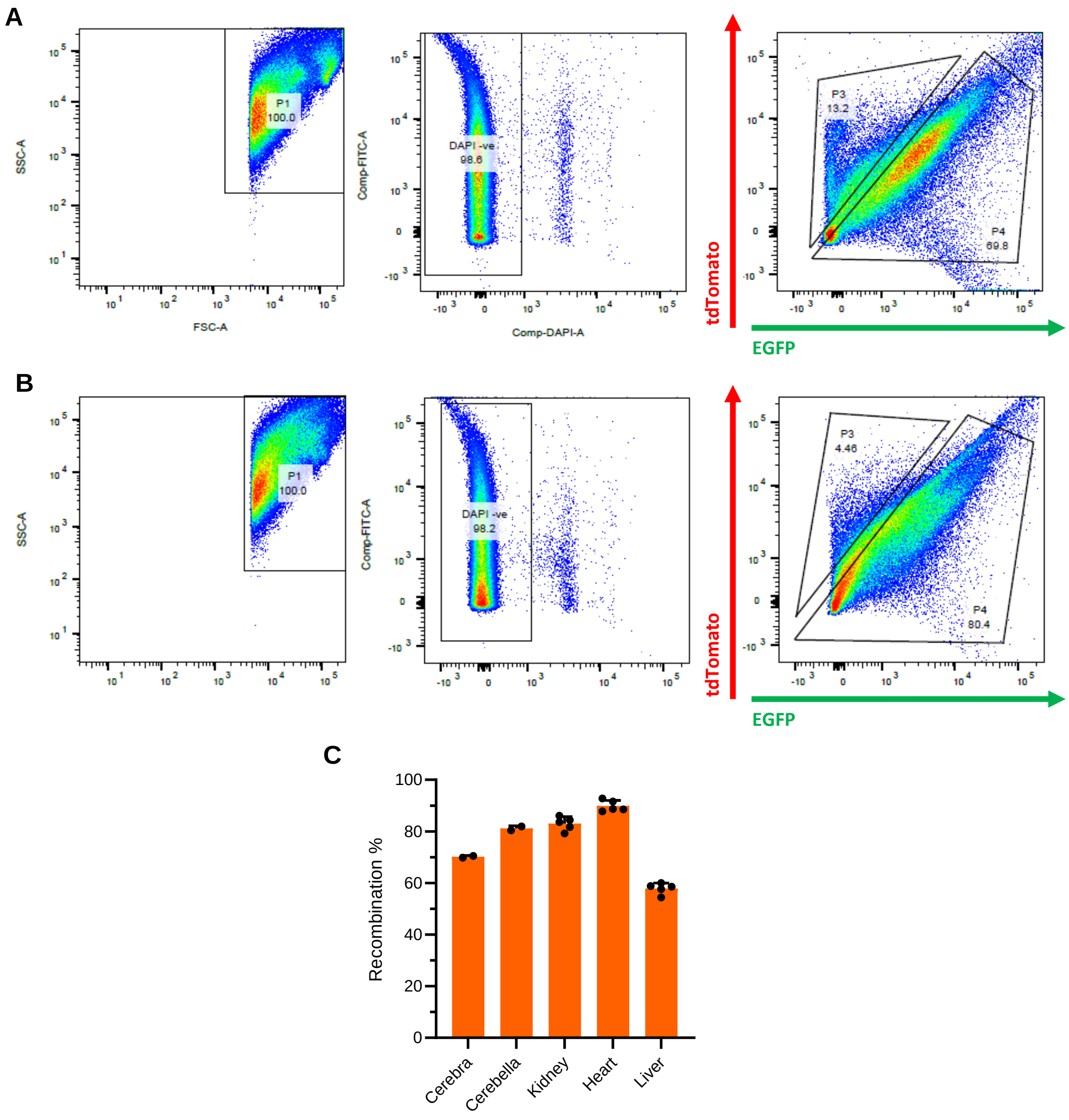


**Supplementary Fig. S1. Analysis of recombination frequency in Polr3a-tamKI mice. (A).** Flow cytometry analysis of a cerebral homogenate from tamoxifen-treated Polr3a-tamKI mice carrying a dual tdTomato-EGFP recombination reporter (Muzumdar et al. 2007). Five tamoxifen injections (6 mg/40 g, i.p) were administered every other day for five days starting at P28. Mice were sacrificed at P42. The plots show side scatter (SSC) and forward scatter (FSC) for area (A) of individual cells, DAPI staining for viability and gating of tdTomato and EGFP based on single color controls. **(B).** Flow cytometry analysis of a cerebellar homogenate from the same mouse as in panel A. **(C).** Recombination frequencies in different tissues. Data for cerebra and cerebella were determined by flow cytometry of two mice treated as described in panel A. Data for kidney, heart and liver were determined by fluorescence microscopy with manual counting of tdTomato and EGFP positive cells. Cell counts were performed on multiple sections with >150 cells scored per section. Results show the mean ± SD.

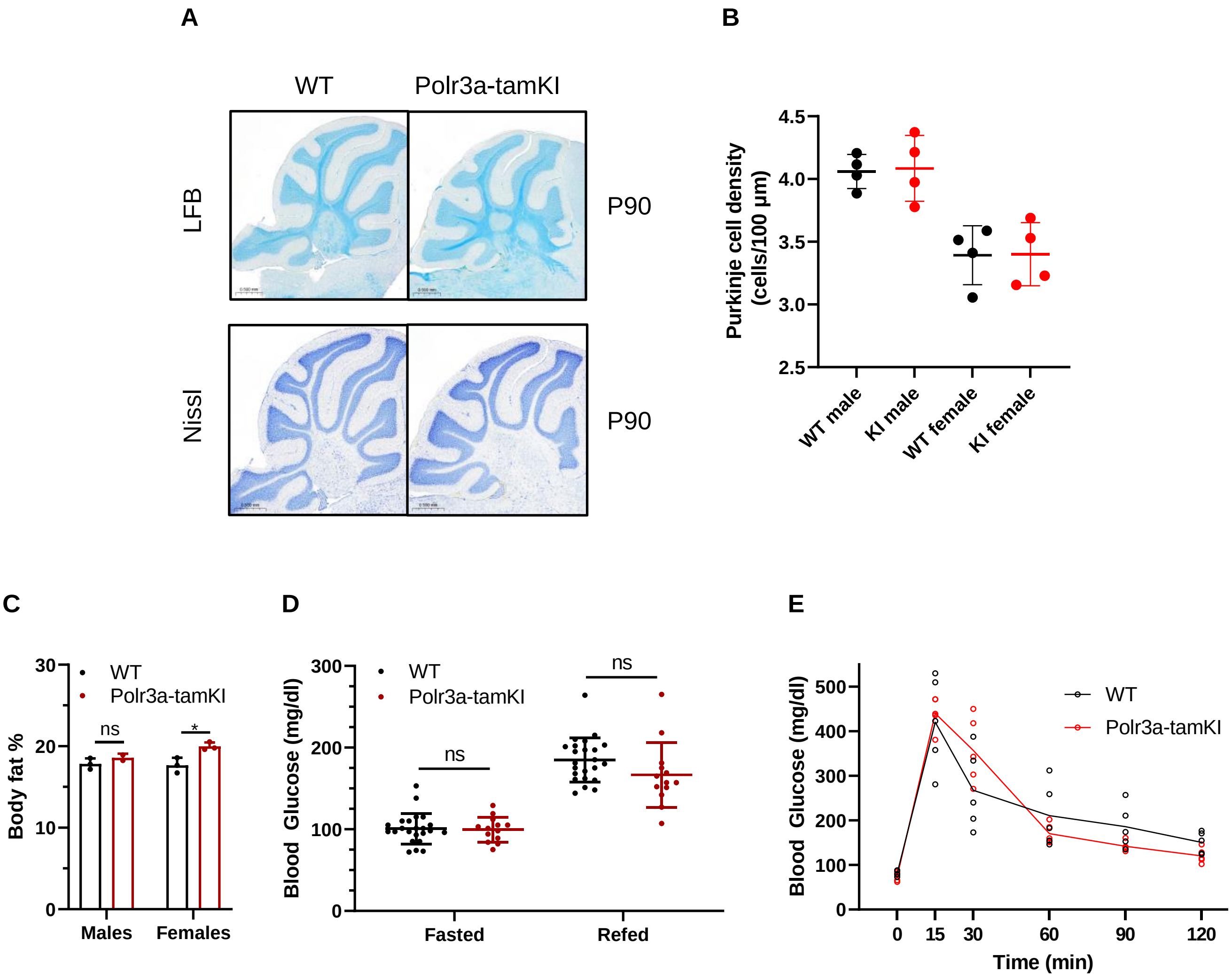


**Supplementary Fig. S2. Cerebellar histology and glucose homeostasis of Polr3a-tamKI mice. (A).** LFB- and Nissl-stained sagittal sections of cerebellum from P90 mice. No differences in staining intensity were noted. **(B)**. Purkinje cell density was quantified in Nissl-stained sections for lobes III, IV/V, VI/VII and VIII. The results are plotted as the mean ± SD for male and female mice of each genotype. **(C)**. Percent body fat was determined by EchoMRI on mice at P42 after an 8 hour midnight fast and a 2 hour refeed. **(D)**. Blood glucose levels (mean ± SD) at P42 were measured by tail vein bleed after an 8-hour midnight fast, and after a 2-hour refeed (WT n=23, Polr3a-tamKI n=13, multiple t-tests, one-way ANOVA). **(E).** Glucose tolerance test. Blood glucose was measured after an 8-hour midnight fast and at 15, 30, 60, 90 and 120 minutes after mice were given a bolus of glucose at 2 g/kg body weight. The lines connect the mean values at each time point. WT and Polr3a-tamKI n=5, multiple t-tests, one-way ANOVA. ns: not significant.

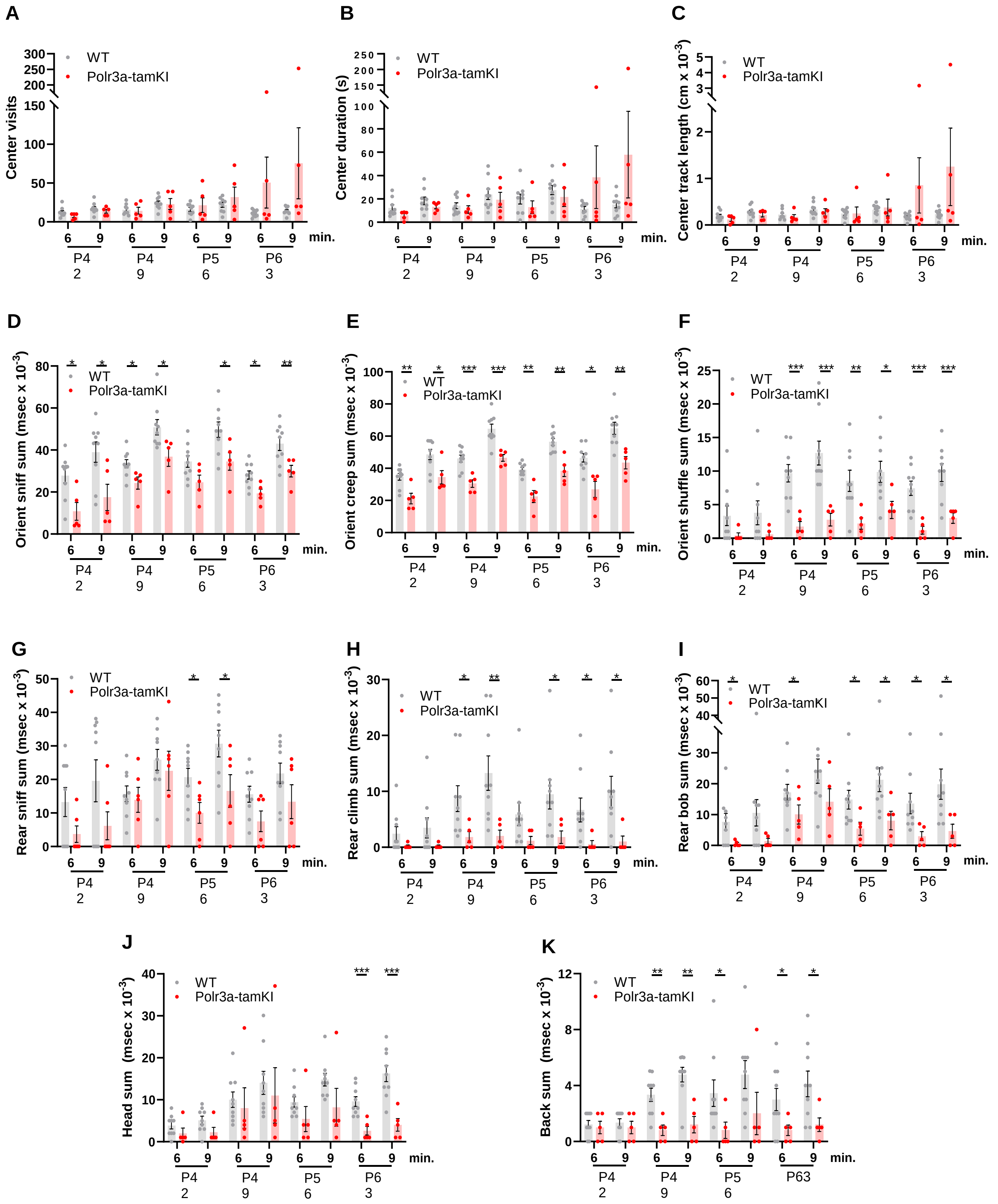


**Supplementary Fig. S3. Behavioral spectrometer analysis of WT and Polr3a-tamKI mice.**

**Supplementary Fig. S3. Behavioral spectrometer analysis of WT and Polr3a-tamKI mice. (A).** Center visit counts. **(B).** Center visit duration. **(C).** Center track length. Anxiety-like behaviors in panels **(A-C)** were scored in a 15 $cm^2$ area in the center of the 40 $cm^2$ arena. **(D).** Orient sniff. **(E).** Orient creep. **(F).** Orient shuffle. Risk assessment behavior was monitored in panels **(D-F)** using three independent metrics. The sum of these behaviors provides an overall measure of risk assessment (see Fig. 3D). **(G).** Rear sniff. **(H).** Rear climb. **(I).** Rear bob. Exploratory behavior was monitored in panels **(G-I)** using three independent metrics. The sum of these behaviors provides an overall measure of exploration (see Fig. 3E). **(J).** Head grooming. **(K).** Back grooming. Data were collected at weekly intervals beginning at P42 (WT n=9 and Polr3a-tamKI n=5). Mice were tested for nine minutes with recording in three intervals of three minutes each. Cumulative data are presented at the 6 and 9 minute time points as the mean ±. SEM. Multiple t-tests and one-way ANOVA, * $p < 0.05$, ** $p < 0.01$, *** $p < 0.001$.

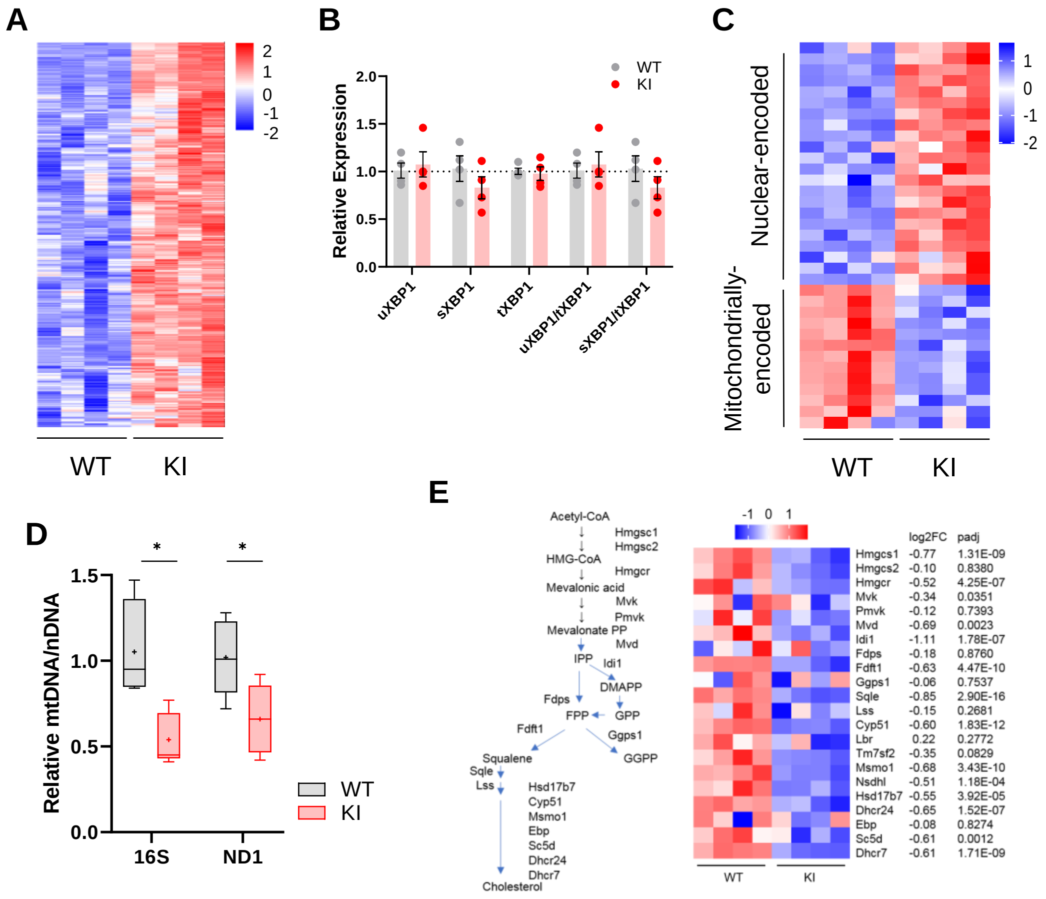


**Supplementary Fig. S4. Gene expression in WT and Polr3a-tamKI (KI) cerebra at P75. (A).** Heatmap represents Z scores of DESeq2 normalized read counts for ATF4-dependent ISR genes activated in cerebra at P75 (p-adj <0.05). **(B).** RT-qPCR analysis of XBP1 splicing. The relative expression of unspliced XBP1 (XBP1u, constitutively expressed) and spliced XBP1 (XBP1s, spliced in response to ER stress) is normalized to total XBP1 mRNA (Yoon et al, 2019). The fold change in expression was calculated by the ΔΔCt method and expressed relative to the WT value. Data are presented as mean ± SEM, n=4 biological replicates. **(C).** A heatmap of mitochondrial and nuclear-encoded mitochondrial proteins showing differential expression (DE) of their genes in KI and WT cerebra at P75. DE genes (p-adj <0.05, log2FC|0.58|) were queried against the Mitocarta 3.0 gene list of mitochondrially-located proteins (Rath et al. 2021) (34/1140 genes). The heatmap represents Z scores of DESeq2 normalized read counts for DE nuclear-encoded and mitochondrially-encoded protein-coding genes (p-adj <0.05, log2FC >|0.58|). **(D).** Relative mitochondrial genome abundance in KI and WT cerebra at P75. PCR analysis of mitochondrial 16S rDNA and ND1 gene abundance was normalized to a nuclear gene (HK2). Fold change was calculated by the ΔΔCt method (Quiros et al. 2017) and expressed relative to the WT value. Box plot and whiskers (min to max) are shown for 16S rDNA and ND1 for WT (gray) and Polr3a-tamKI (red), 3-4 biological replicates repeated 3 times. Median is represented by +, * p ≤ 0.05 (Student's standard t-test). **(E).** Sterol biosynthesis is down-regulated in KI cerebra. Scheme of sterol biosynthesis (adapted from Rye et al. 2018) and gene expression changes of enzymes annotated to the GO term sterol biosynthetic process pathway for cholesterol synthesis (GO:0016126). The heatmap shows Z-scores of DESeq2 normalized read counts adjacent to the log2FoldChange and p-adj values from Supplementary Table S1.

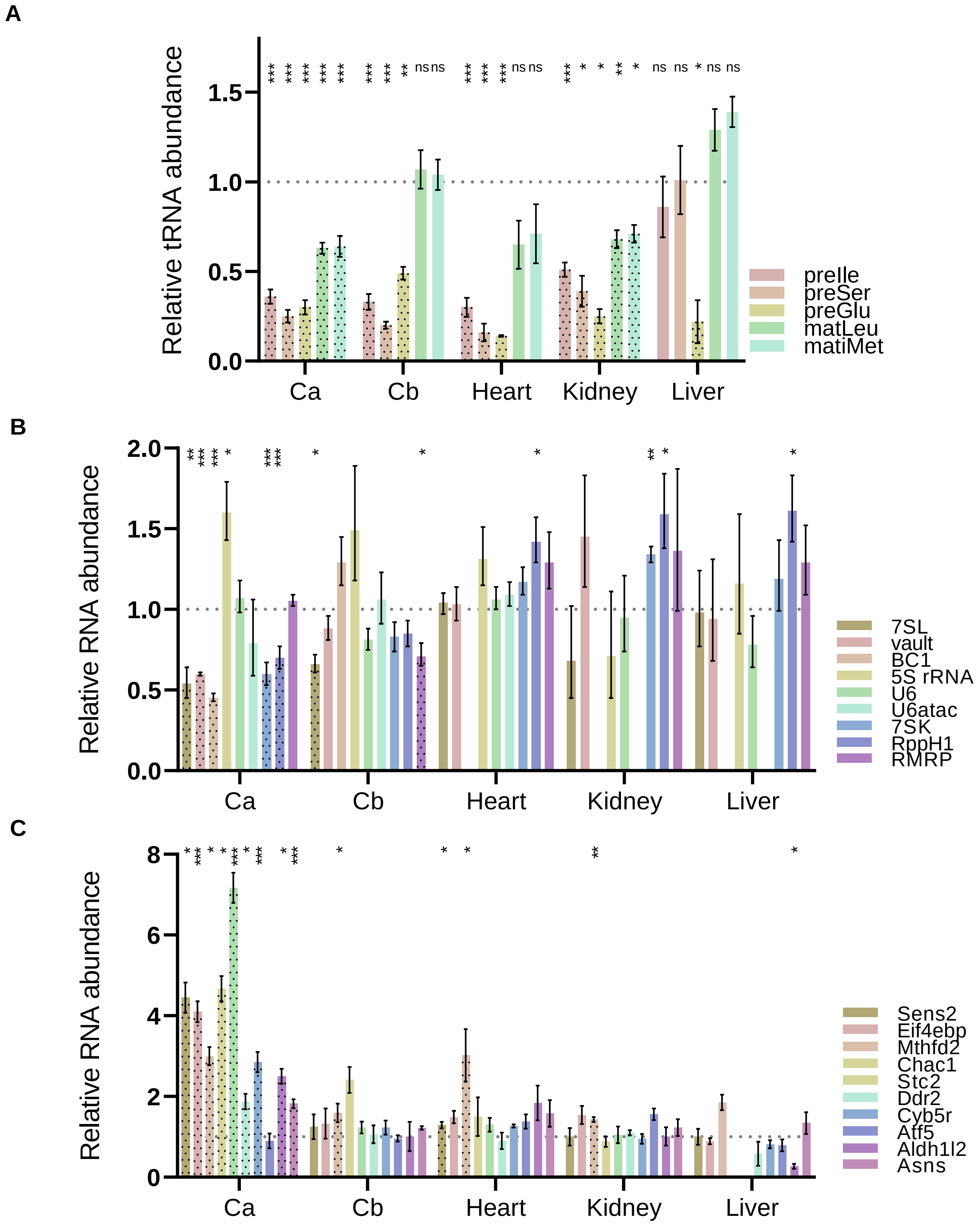


**Supplementary Fig. S5. Pol III and Pol II transcript levels in various tissues from WT and Polr3a-tamKI (KI) mice at P75.**

**Supplementary Fig. S5. Pol III and Pol II transcript levels in various tissues from WT and Polr3a-tamKI (KI) mice at P75. (A).** The relative abundance of precursor and mature tRNAs in total RNA from cerebella (Cb), heart, kidney and liver. Cerebral (Ca) data (from Figs. 3B,C) are included for ease of comparison. Precursor-tRNAIle-TAT (preIle) and mature tRNAs Leu-CAA and iMet (matLeu and matiMet, respectively) levels were determined by northern blotting of 5 to 6 biological replicates and normalized to a U3 snRNA loading control. RT-qPCR of 4 to 6 biological replicates was used to determine relative pre-tRNA levels for pre-tRNASer *Ts12* and pre-tRNAGlu *Te5/7/8* (preSer and preGlu, respectively). The fold change in expression was calculated by the $\Delta\Delta$Ct method (Taylor et al. 2019) and expressed relative to the WT value. Data represent the mean ± SEM. The results for preIle, preSer, preGlu, matLeu and matiMet are plotted from left to right for each tissue. Only KI values are shown for simplicity. tRNAs with significant differences between WT and Polr3a-tamKI samples are highlighted with black dots. ns, not significant; * $p \leq 0.05$; ** $p \leq 0.01$; *** $p \leq 0.005$ (Student's standard t-test). **(B).** RT-qPCR analysis of non-tRNA Pol III transcripts across WT and KI cerebella (Cb), heart, kidney and liver are shown. Cerebral (Ca) values (from Fig. 3C) are included for ease of comparison. Only KI values are shown for simplicity. Pol III transcripts with significant differences between WT and KI are highlighted with black dots. Data were calculated, normalized and plotted as in panel **A**, n= 3-6 biological replicates. **(C).** RT-qPCR analysis of a set of ATF4-regulated ISR genes across WT and KI cerebella (Cb), heart, kidney and liver are shown. Cerebral (Ca) values (from Fig. 3D) are included for ease of comparison. Data were calculated, normalized and plotted as in Fig. 3D, n= 3-5 biological replicates.

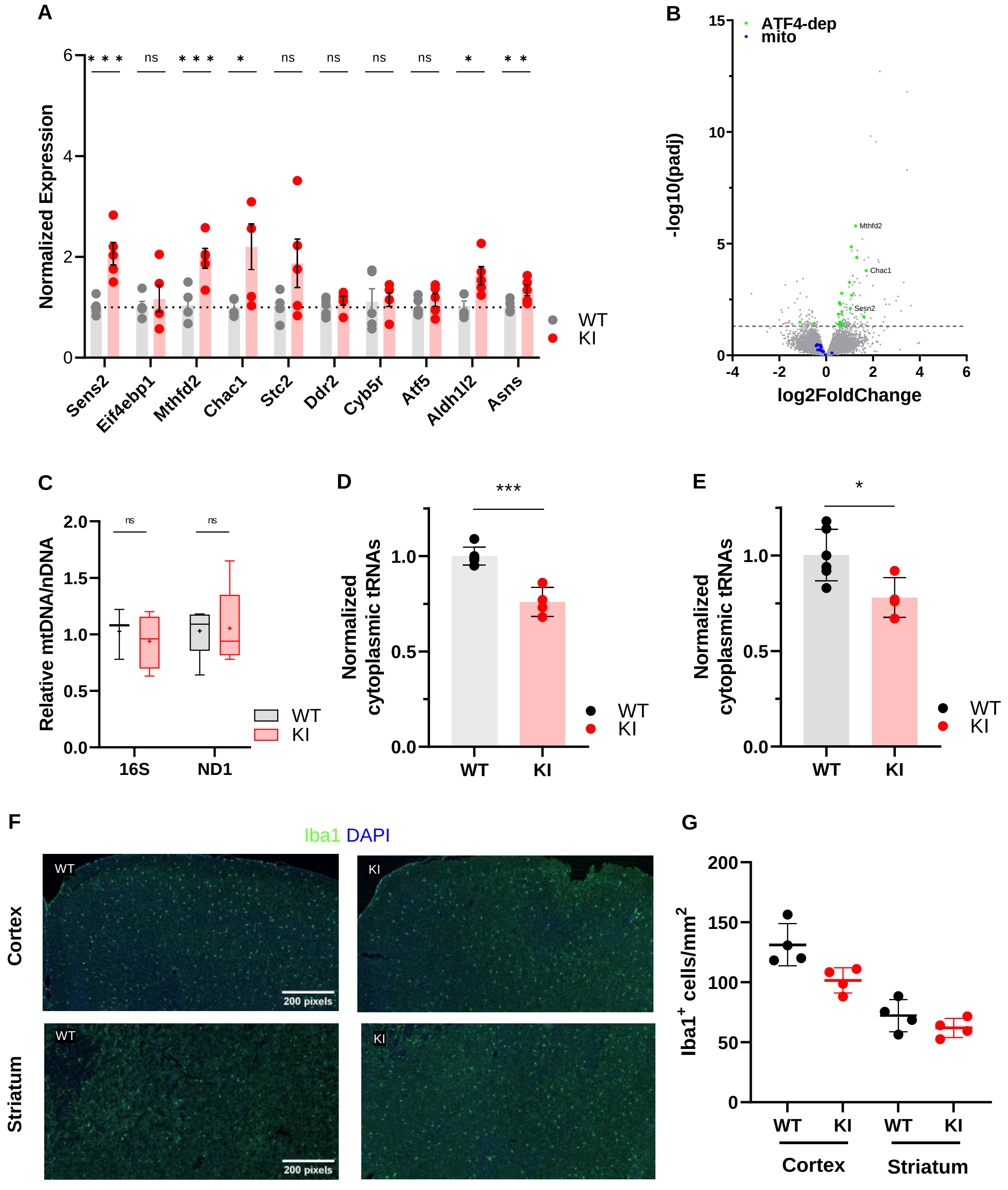


**Supplementary Fig. S6. Gene expression and Iba1 staining in adolescent WT and Polr3a-tamKI (KI) cerebra.**

**Supplementary Fig. S6. Gene expression and Iba1 staining in adolescent WT and Polr3a-tamKI (KI) cerebra. (A).** RT-qPCR analysis of a set of ATF4-regulated ISR genes in WT and KI cerebra at P42. Data were calculated, normalized and plotted as in Fig. 3D, n= 3-5 biological replicates. ns, not significant; * $p \leq 0.05$; ** $p \leq 0.01$; *** $p \leq 0.005$ (Student's standard t-test). **(B).** Fold change in global gene expression in WT and KI cerebra at P42. A volcano plot shows RNA-seq expression changes (KI/WT, n=4 biological replicates). Differentially expressed (DE) ATF4-regulated ISR genes are highlighted in green (p-adj <0.05, log2FC >|0.58|). Mitochondrial-encoded mRNAs are shown in blue for comparison with Fig. 3E. Representatives of the ATF4-regulated ISR gene set assayed in panel **A** are labeled. **(C).** Relative mitochondrial genome abundance in KI and WT cerebra at P42. PCR analysis of mitochondrial 16S rDNA and ND1 gene abundance was calculated, normalized and expressed as described in Supplemental Fig. S4D. Box plot and whiskers (min to max) are shown for 16S rDNA and ND1 for WT and KI cerebra at P42, 4-5 biological replicates, replicated 3 times. Median is represented by +, ns, not significant (Student's standard t-test). **(D-E).** Total tRNA reads in WT and KI cerebra at P42 were normalized to the sum of endogenous mitochondrially-encoded tRNA reads **(D)** or exogenous spike-in reads **(E)** and expressed relative to the mean WT value. Data represent the mean ± SD, KI n=4 and WT n=6 biological replicates, p = 0.0003 and 0.024 (Student's standard t-test) for panels **D** and **E**, respectively. **(F).** Iba1 staining of microglia in the cerebral cortex and striatum at P44. Scale bar, 200 µm, applies to all panels. **(G).** Iba1+ cell counts in the cerebral cortex and striatum represent the mean ± SD of bilaterally symmetric regions, (n= 2).

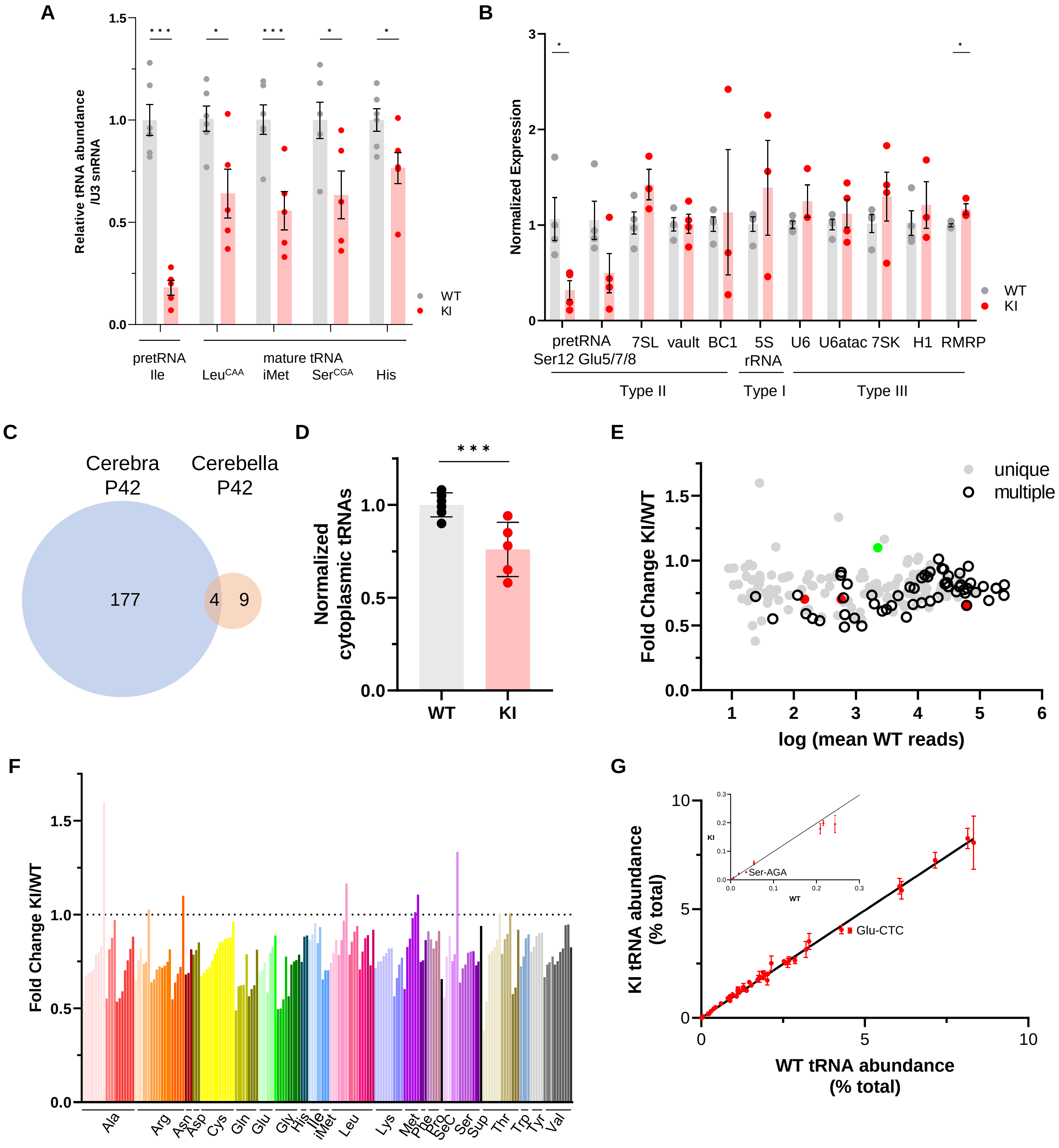


**Supplementary Fig. S7. Gene expression in WT and Polr3a-tamKI (KI) cerebella at P42.**

**Supplementary Fig. S7. Gene expression in WT and Polr3a-tamKI (KI) cerebella at P42. (A-B).** The relative abundance of Pol III transcripts in total RNA from WT and KI cerebella (Cb) at P42 determined by northern analysis **(A)** and RT-qPCR **(B)**. **(A).** Precursor (pre-tRNAIle-TAT) and mature tRNAs (Leu-CAA, iMet, Ser-CGA, His) detected by northern blotting were quantified as in Fig. 3A. Data represent the mean ± SEM, n=6 WT and n=5 KI biological replicates. **(B).** Pol III transcripts detected by RT-qPCR in KI and WT Cb. Data were normalized as in Fig. 3C and represent the mean ± SEM, n= 3-6 biological replicates; ns, not significant; * p ≤0.05; ** p ≤0.01; *** p ≤0.005 (Student's standard t-test). **(C).** Venn diagram shows limited overlap between significant DE genes from cerebra (Ca) and cerebella (Cb) at P42 (p-adj <0.05, log2F|0.58|), KI n=4 and WT n=4 biological replicates for both Ca and Cb. **(D).** Total cytoplasmic tRNA reads in WT and KI cerebella at P42 were normalized to the sum of endogenous mitochondrially-encoded tRNA reads and exogenous spike-in reads and expressed relative to the mean WT value. Data represent the mean ± SD, KI n=5 and WT n=5 biological replicates, p = 0.0006 (Student's standard t-test). **(E).** Fold change for KI/WT cytoplasmic tRNAs is plotted against log mean WT reads. The symbols show tRNAs encoded by unique loci (gray), identical tRNAs encoded by multiple loci (hollow black), iMet tRNAs (red) and tRNAArg-TCT-4-1 (green). **(F).** Fold change for cytoplasmic tRNAs (KI/WT) is plotted for all decoder families. Individual tRNAs are ordered from most to least fold change and grouped by codon recognition (tRNA decoder) family. The amino acid that is charged by each tRNA decoder family is indicated. **(G).** The cytoplasmic tRNA profile for KI P42 cerebella is plotted against the WT profile. tRNA decoder reads from each KI and WT replicate were expressed as a percentage of their respective total cytoplasmic decoder pool. Data represent the mean ± SEM. tRNA decoders that are significantly lower in KI compared to WT (p ≤ 0.05, Student's standard t-test) fall below the regression line and are labeled. The inset shows tRNA decoders with <0.3% of the tRNA pool.
